# Autoregulation of RNase E synthesis: Predominant impact of a branched RNA stem-loop to which the nuclease binds

**DOI:** 10.64898/2026.10.07.757400

**Authors:** Salah Eldein Elkattawy, Tithi Banerjee, Joel G. Belasco

## Abstract

Due to its crucial role in proteobacterial RNA processing and degradation, the synthesis of the endonuclease RNase E is tightly controlled by a post-transcriptional autoregulatory mechanism governed by the 5′ untranslated region of *rne* mRNA, the transcript that encodes it. Previous studies have identified one structural component of the *rne* 5′ untranslated region (hp2) that helps to mediate this process in *Escherichia coli* by binding to RNase E. Here, we show that another component (hp3), a branched stem-loop, makes an even greater contribution to feedback regulation of RNase E synthesis and that the effects of hp2 and hp3 are additive, suggesting independent function. Despite the structural complexity of hp3, a single constituent branch appears to be the principal source of its regulatory activity. This branch also enables hp3 to bind RNase E with high affinity. The secondary structure of this branch, including the presence of two internal loops essental for its impact, is conserved in diverse Enterobacterales species, whereas the sequence of the internal loops is not. We propose that hp2 and hp3 mediate autoregulation by recruiting RNase E to the *rne* transcript.

## INTRODUCTION

Messenger RNA (mRNA) degradation is a mechanism of great importance for controlling gene expression throughout biology due to its direct impact on the number of times that a given message can be translated by ribosomes. The half-lives of the various mRNAs in a cell can differ substantially from one another, ranging from seconds to about an hour in bacteria and from minutes to days in humans. Moreover, like rates of transcription and translation, mRNA lifetimes can be modulated in response to a cell’s needs.

By far, the most important ribonuclease for controlling RNA lifetimes in proteobacteria is the endonuclease RNase E, which not only triggers the degradation of most mRNAs and many noncoding small RNAs (sRNA) but also participates in the maturation of ribosomal RNA (rRNA) and transfer RNA (tRNA) from precursor transcripts (1). RNase E cuts RNA internally with only modest sequence specificity in unpaired regions that are AU-rich (2,3). The rate at which it cleaves mRNA is governed primarily by other characteristics of the transcript, such as ribosome binding/translation, protein binding, and RNA structure, as well as the phosphorylation state of the 5′ terminus, where a 5′ monophosphate can act to recruit RNase E (4–10).

In view of its important role in RNA processing and degradation, it is not surprising that RNase E is an essential enzyme in proteobacteria. In fact, not only the underproduction of this enzyme but also its overproduction is deleterious to cell growth (11–13). To keep the cellular concentration of RNase E within a narrow range, *E. coli* has evolved an autoregulatory mechanism in which RNase E degrades its own mRNA (the transcript of the *rne* gene) at a rate that is unusually sensitive to the cellular level of RNase E activity (14,15). Feedback regulation of *rne* gene expression in *E. coli* is mediated in cis by the 361-nt *rne* 5′ untranslated region (UTR), which can confer this property onto other mRNAs (e.g., *lacZ*) to which it is translationally fused in-frame (15). Previous research in *E. coli* has demonstrated an important regulatory role for one stem-loop in particular within the *rne* 5′ UTR (hp2) and has hinted at a smaller role for another, more complex base-paired structure there (hp3) (16). Additional studies have mapped the influence of hp2 on feedback regulation to the upper one-third of this stem-loop, a region comprising a 4-nt apical loop and an 8-nt internal loop separated by two base pairs, and have shown that it serves as a binding site for RNase E (17,18). By contrast, almost nothing is known about the contribution of hp3 and its various components to feedback regulation.

We have now quantified the relative impact of *rne* hp3 on the autoregulation of RNase E synthesis and identified the regions of this branched stem-loop that are responsible for its influence. These studies show that the contribution of hp3 to feedback regulation is greater than that of hp2 and that their effects are roughly additive. In addition, these experiments reveal that the impact of hp3 on this important regulatory process is primarily attributable to a single branch of this complex base-paired structure and that this branch enables hp3 to bind RNase E with high affinity.

## RESULTS

Despite extensive sequence divergence (∼70%), the secondary structure of the 5′ UTR is well conserved in the *rne* transcripts of diverse Enterobacterales, an order of Gram-negative bacteria of which *E. coli* and many other pathogens are members (Figures 1 and S1). It comprises three stem-loops (hp1, hp2, and hp3) that are separated from one another by short unpaired RNA segments and followed by a third unpaired region containing the signals for translation initiation (Figure 2A). The structure as diagrammed is consistent with the results obtained by subjecting the *E. coli rne* transcript to chemical modification *in vitro* and *in vivo* (16). Previous research has shown that removing a large portion of the *rne* 5′ UTR encompassing both hp2 and hp3 virtually abolishes feedback regulation of RNase E synthesis in *E. coli* and that feedback regulation is partially impaired by deleting smaller regions encompassing either hp2 or hp3 individually, whereas deleting hp1 has little or no effect (16). However, the design of those experiments had three significant flaws. First, the deletions often removed not only the stem-loop but also adjacent RNA segments. Second, deletions of this kind directly join nucleotides that ordinarily are spatially separated from one another, thereby altering the distance between flanking RNA elements. Third, by completely excising each of these stem-loops, such deletions may disrupt the tertiary structure of the *rne* 5′ UTR by preventing other stem-loops from stacking on hp2 and hp3. We therefore adopted an improved strategy for examining the contributions of hp2 and hp3 to feedback regulation by instead truncating these stem-loops, individually or in combination (Figure 2B). The effect of these truncations in the context of a plasmid-borne *rne-lacZ* reporter was then examined by comparing reporter gene expression in *E. coli* cells containing a low or high level of RNase E activity.

**Figure 1.**
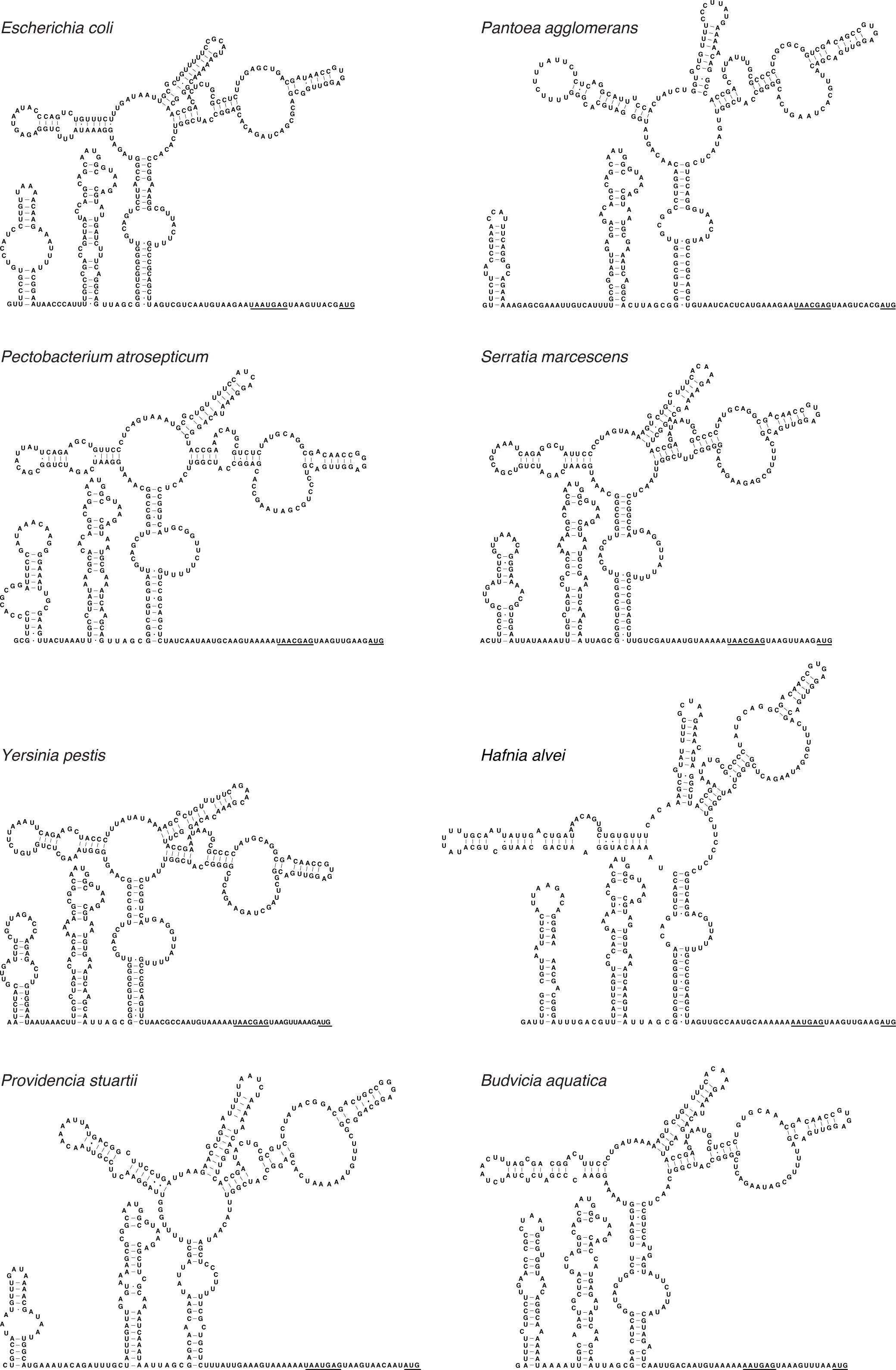
Conserved secondary structure of the *rne* 5′ UTR. The phylogenetically conserved secondary structure of the 5′ UTR is depicted for *rne* transcripts from eight proteobacterial species representing seven families of the order Enterobacterales: Enterobacteriaceae (*Escherichia coli*), Erwiniaceae (*Pantoea agglomerans*), Pectobacteriaceae (*Pectobacterium atrosepticum*), Yersiniaceae (*Serratia marcescens* and *Yersinia pestis*), Hafniaceae (*Hafnia alvei*), Morganellaceae (*Providencia stuartii*), and Budviciaceae (*Budvicia aquatica*). The 5′ end of the *E. coli* transcript was previously mapped by primer extension analysis (15), while that of the other transcripts was deduced from the location of the *rne* promoter, whose sequence is conserved in these eight species (TTGCTA – 18 bp – AATAAT, except TTGCTA – 18 bp – GATAAT in *Providencia stuartii* and TTGCCA – 17 bp – GATAAT in *Budvicia aquatica*). The Shine-Dalgarno element and initiation codon of each transcript are underlined.

**Figure 2.**
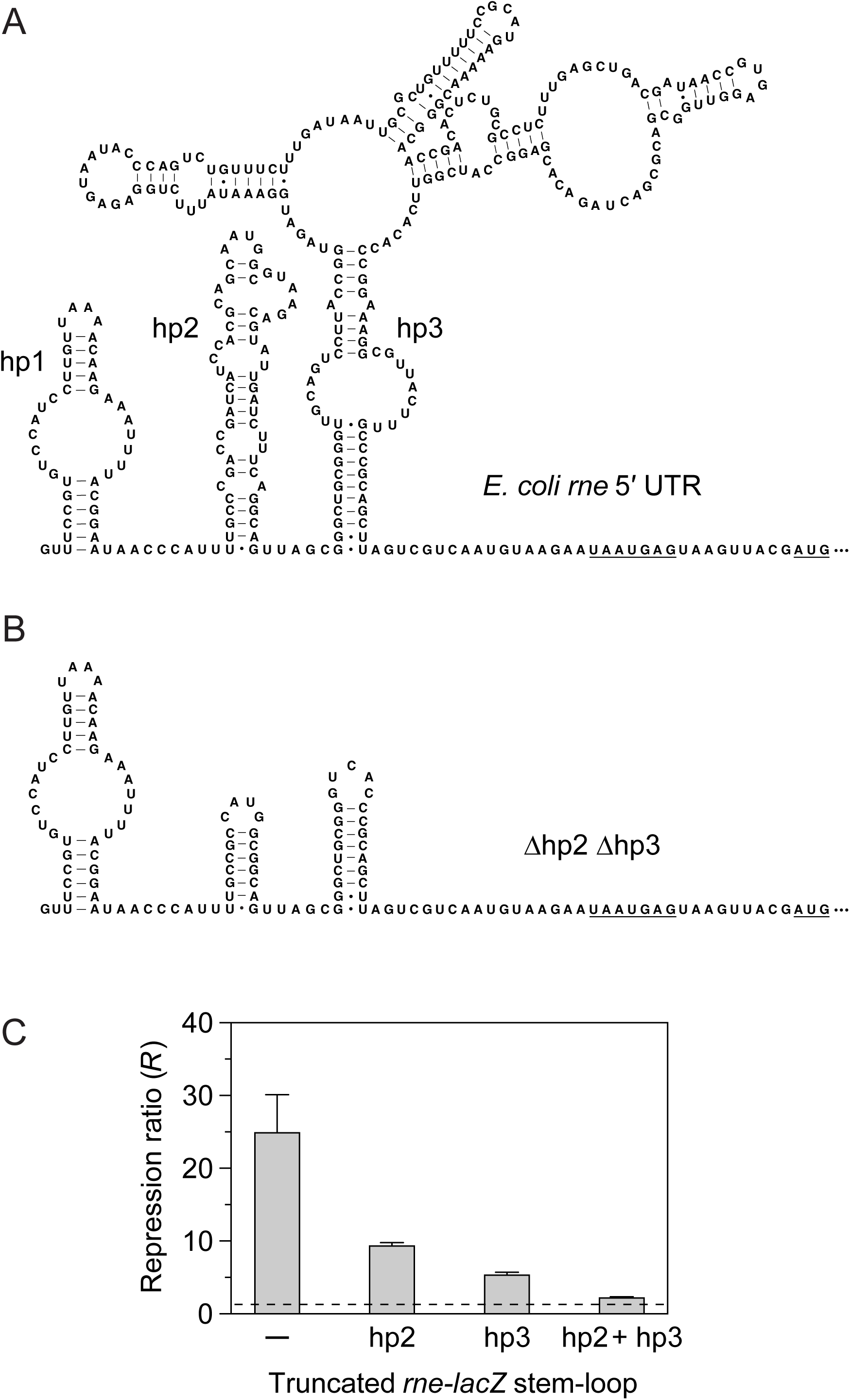
*rne* 5′ UTR elements that mediate the autoregulation of RNase E synthesis. (A) Secondary structure of the *E. coli rne* 5′ UTR. Three base-paired structures within the *rne* 5′ UTR (hp1, hp2, and hp3) are labeled, and the Shine-Dalgarno element and initiation codon are underlined. (B) Truncation of *rne* hp2 and hp3. A mutated *rne* 5′ UTR in which both hp2 and hp3 have been truncated (Δhp2 Δhp3, encoded by pEZ1223) is shown. In other mutants (not shown), only one (Δhp2, encoded by pEZ1202) or the other (Δhp3, encoded by pEZ1203) of these structures was truncated. (C) Repression ratios. The ratio of β-galactosidase produced from an *rne-lacZ* reporter in *E. coli* cells containing a low versus a high level of RNase E activity (*R*) is shown for the wild-type *rne* 5′ UTR (–) and for three mutants in which hp2 and/or hp3 had been truncated. The dashed line corresponds to a repression ratio of 1 (no regulation).

A low level of cellular RNase E activity was achieved by using *E. coli* strain CH1828 as the host and growing the cells to log phase at 37°C. At this semi-permissive temperature, the mutant form of RNase E encoded by the temperature-sensitive *rne*-1 (*ams*-1) allele of CH1828 is only partially active (15). A high level of cellular RNase E activity was achieved by overproducing wild-type RNase E 2.8-fold in an isogenic *rne*^+^ strain, CH1827, containing a multicopy plasmid (pRNE101) that encodes the wild-type enzyme (15). The expression of a plasmid-encoded *rne-lacZ* reporter and its mutant countparts bearing truncations of hp2 and/or hp3 in the *rne* 5′ UTR was quantified in each of these strains by assaying for β-galactosidase activity.

As a consequence of feedback regulation, the level of β-galactosidase activity observed for the wild-type *rne-lacZ* reporter encoded by plasmid pEZ1200 was 25-fold greater (repression ratio (*R*) = 24.97 ± 5.13) in cells containing a low level of RNase E activity versus a high level of RNase E activity (Figure 2C). Truncating either stem-loop substantially reduced *rne-lacZ* regulation, resulting in diminished repression ratios of 9.43 ± 0.36 in the absence of hp2 (pEZ1202) and 5.41 ± 0.32 in the absence of hp3 (pEZ1203). The repression ratio was even smaller when both hp2 and hp3 were truncated (pEZ1223), which lowered *R* to a value (2.27 ± 0.10) barely above the minimum (1.00). We conclude that both hp2 and hp3 contribute to feedback regulation, that their effects are roughly additive, and that feedback regulation is nearly abolished when both are truncated. Interestingly, these measurements also show that hp3 has almost twice the impact of hp2, contradicting a previous assertion that hp2 predominates (16).

Due to its great importance for feedback regulation, we next dissected hp3 to determine the source of its regulatory influence. This base-paired structure is a branched stem-loop that comprises a four-way junction connecting stems that each contain one or more internal loops or bulges (Figure 2A). We designated its four component stems and stem-loops hp3A, hp3B, hp3C, and hp3D and its central hub hp3H (Figure 3A). First, stem-loops hp3B, hp3C, and hp3D were each truncated individually so as to shorten them, remove any internal loops or bulges, change the sequence of the base-paired stem, and replace the apical loop with a GAAA tetraloop. Separately, the large internal loop of hp3A was removed. The regulatory impact of each of these mutations (Δhp3A, Δhp3B, Δhp3C, and Δhp3D (Figure 3A)) and every subsequent hp3 mutation was then examined in the context of an *rne-lacZ* reporter lacking hp2 (i.e., in a derivative of pEZ1202, for which *R* = 9.43 ± 0.36) so as not to obscure the effects of the hp3 mutations. (For the sake of brevity, the absence of hp2 has been omitted from the names of these hp3 mutants.) The repression ratios measured for the four new hp3 mutants showed that truncating hp3D caused by far the greatest reduction of *R* (3.45 ± 0.38), followed by truncating hp3A (6.00 ± 0.98), hp3C (6.90 ± 0.81), and hp3B (8.57 ± 1.40), the last of which had no detectable effect on hp3 function (*p* = 0.94 versus 9.43 ± 0.36 for pEZ1202).

**Figure 3.**
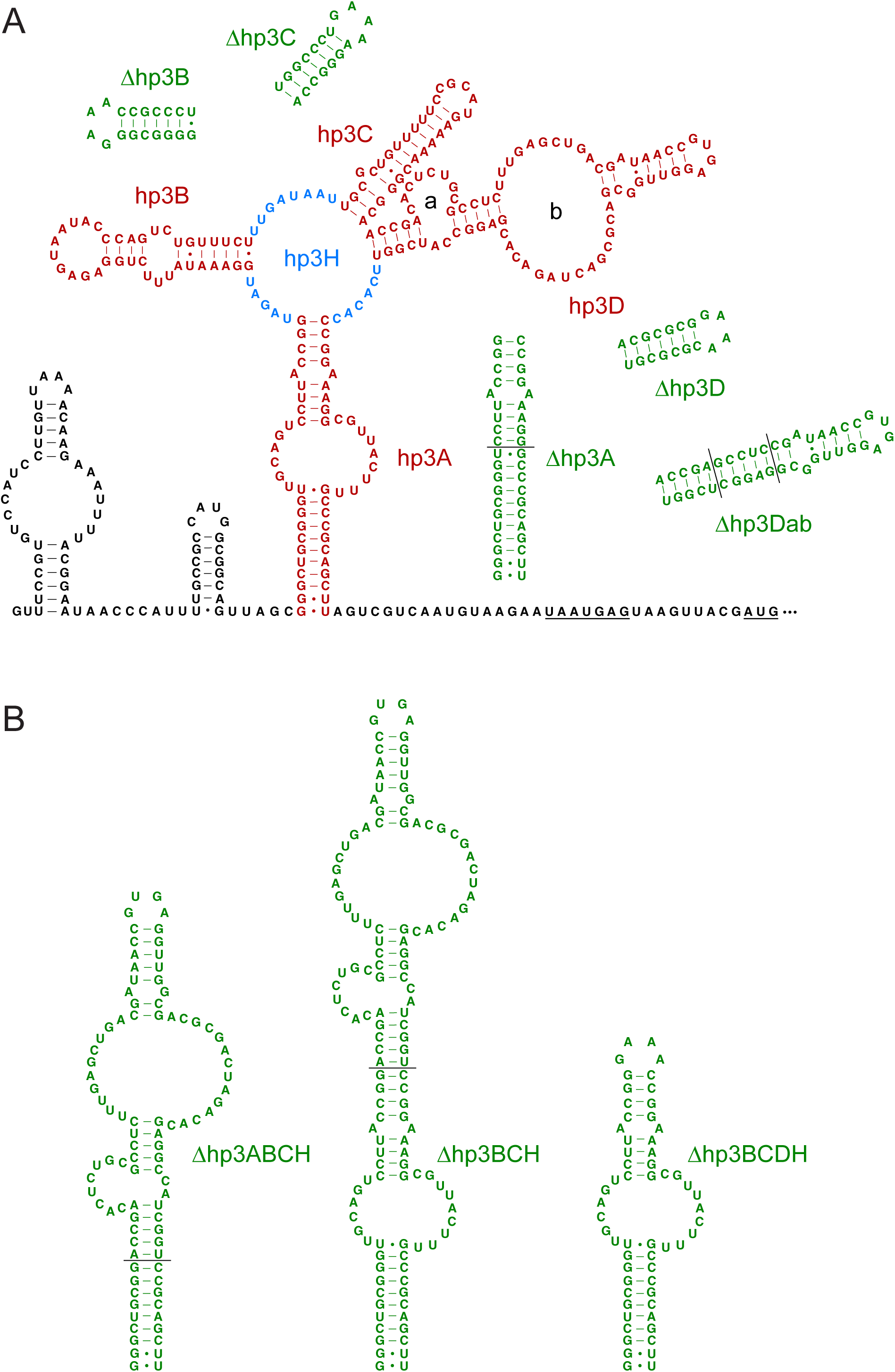
*rne* hp3 deletion mutants. (A) Deletion of hp3 components from the *rne* 5′ UTR. The structural components of *rne* hp3 are depicted in the context of a pEZ1202-encoded *rne-lacZ* reporter lacking hp2, with hp3A, hp3B, hp3C, and hp3D colored red and the four-way junction (hp3H) colored blue. In addition, the two internal loops of hp3D (a and b) are labeled. Shown in green beside the four red components of hp3 are truncation mutants of each (Δhp3A, Δhp3B, Δhp3C, and Δhp3D) as well as a mutated variant of hp3D lacking both internal loops (Δhp3Dab). In other hp3D mutants (Δhp3Da and Δhp3Db, not shown), only one or the other internal loop was removed. (B) *rne* hp3 mutants lacking the four-way junction. Three mutated variants of hp3 lacking hp3B, hp3C, and the four-way junction (hp3H) are shown. In Δhp3ABCH and Δhp3BCH, hp3D was mounted directly atop hp3A, with or without removing the large internal loop and the upper half of hp3A. In Δhp3BCDH, all of hp3 except hp3A was removed.

To maximize reproducibility and enable accurate quantitative comparisons of data from experiments performed on different days, the repression ratios of the progenitor pEZ1202-encoded reporter in which hp3 remained intact and of the pEZ1223-encoded reporter in which hp3 had been truncated (hereafter referred to as Δhp3) were re-measured every time that a group of hp3 mutants was tested. The fold effect of deleting each hp3 component (*ED*) was then calculated by dividing the repression ratio measured for the progenitor pEZ1202 reporter by that of each hp3 mutant measured in parallel after first subtracting 1 from each value because a repression ratio of 1 indicates no feedback regulation whatsoever: *ED* = (*R*_intact hp3_ –1) / (*R*_mutant hp3_ –1), where *ED* ≥ 1 (the limiting value for a mutation with no effect). These calculations revealed that deleting hp3D had more than a three-fold effect on feedback regulation (*ED* = 3.12 ± 0.71), whereas deleting hp3C or the internal loop of hp3A had only a 1.3- or 1.5-fold effect, respectively, and deleting hp3B had no detectable effect (Figure 4A, Table S1). For comparison, deleting hp3 in its entirety (Δhp3) had a 7.22 ± 0.67 fold effect.

**Figure 4.**
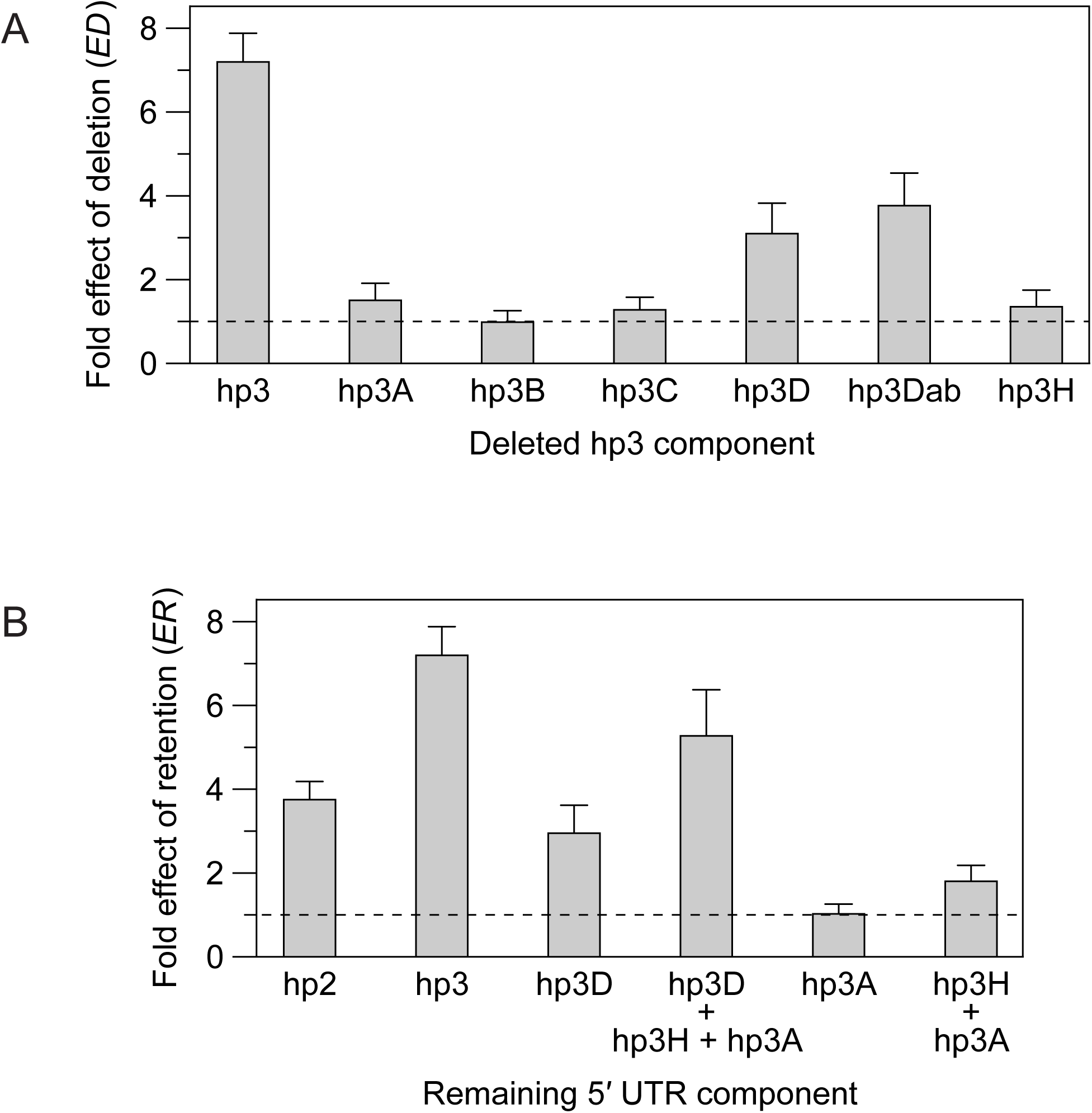
Regulatory impact of elements within the *rne* 5′ UTR. (A) Effect of deleting hp3 components. The fold effect (*ED*) on *rne-lacZ* expression of removing *rne* hp3 or various components thereof was examined in a reporter lacking hp2. The dashed line corresponds to a one-fold effect (i.e., no effect of the deletion whatsoever). (B) Sufficiency of individual *rne* 5′ UTR components for feedback regulation. The fold effect (*ER*) on *rne-lacZ* expression of retaining various components of *rne* hp3 in the absence of any others was examined in a reporter lacking hp2. For comparison, the fold effect of the presence of either hp2 or intact hp3 is also shown. The dashed line corresponds to a one-fold effect (i.e., no effect of the retained 5′ UTR element whatsoever).

Stem-loop hp3D contains two large internal loops. They were each deleted individually (Δhp3Da or Δhp3Db) or in combination (Δhp3Dab) to determine their contributions to feedback regulation (Figure 3A). The combined effect of removing both internal loops (*ED* = 3.79 ± 0.76 for Δhp3Dab) was similar to the effect of deleting hp3D (3.12 ± 0.71 for Δhp3D; *p* = 0.34) and greater than the effect of removing either internal loop individually (*ED* = 2.04 ± 0.49 and 2.63 ± 0.53 for Δhp3Da and Δhp3Db, respectively) (Figure 4A). We conclude that hp3D relies on its two large internal loops for its regulatory activity.

To assess the regulatory contribution of the central hub of hp3 (hp3H), we first measured the repression ratio of a mutant in which stem-loops hp3B and hp3C had both been truncated and the large internal loop of hp3A had been removed (Figure 3A). As expected from the minor effect of mutating each of these hp3 components individually, the effect of mutating all three together (Δhp3ABC, *ED* = 1.75 ± 0.34) was small. We then examined the effect of additionally deleting the central hub (hp3H). This was accomplished by mounting hp3D directly atop the lower portion of hp3A (Figure 3B). The fold effect of combining all four mutations (Δhp3ABCH, *ED* = 2.42 ± 0.47) was only marginally greater than that measured for Δhp3ABC. A direct comparison of the two mutants (*ED*_Δhp3ABCH_/*ED*_Δhp3ABC_) revealed that the regulatory defect caused by deleting hp3H was only 1.38 ± 0.38 fold (Figure 4A). This finding indicates that the four-way junction of hp3 makes only a small contribution to the impact of this branched stem-loop on feedback regulation.

Because the contribution of the large internal loop of hp3A to feedback regulation is not negligible, we also compared two other mutants in which only stem-loops hp3B and hp3C had been truncated (Δhp3BC) and in which hp3H had additionally been removed so that only hp3D and hp3A remained (Δhp3BCH) (Figure 3B). The fold effect of deleting hp3H calculated by comparing these two mutants (*ED*_Δhp3BCH_/*ED*_Δhp3BC_, Table S1) was 1.48 ± 0.42, a value similar to that calculated by comparing the corresponding mutants from which the large internal loop of hp3A had also been deleted (1.38 ± 0.38; see above). These findings confirm the relatively modest contribution of the four-way junction to the regulatory impact of hp3.

By contrast, the retention of hp3D, hp3H, and hp3A in combination is sufficient to mediate feedback regulation with nearly three-quarters of the efficacy of retaining hp3 in its entirety, as evidenced by the fold effect of retention (*ER*) calculated by comparing the Δhp3BC mutant to the Δhp3 mutant (*ER* = (*R* _Δhp3BC_ –1) / (*R* _Δhp3_ –1) = *ED*_Δhp3_/*ED*_Δhp3BC_ = 5.30 ± 1.08 versus 7.22 ± 0.67 when hp3 is left intact) (Figure 4B, Table S2). The fold effect of retaining hp3D alone (*ER* = 2.98 ± 0.64), determined by comparing the Δhp3ABCH and Δhp3 mutants (*ED*_Δhp3_/*ED*_Δhp3ABCH_), was also substantial: almost half the effect of retaining hp3 in its entirety and nearly as large as the effect of retaining only hp2 (*ER* = 3.78 ± 0.41) (Figure 4B, Table S2). Conversely, retaining hp3H and hp3A in combination had a relatively small effect (*ER* = *ED*_Δhp3_/*ED*_Δhp3BCD_ = 1.82 ± 0.36), and retaining hp3A alone was not sufficient to significantly regulate *rne-lacZ* expression, as the fold effect observed for an hp3 mutant in which only hp3A remained intact (Δhp3_BCDH_, *ER* = *ED*_Δhp3_/*ED*_Δhp3BCDH_) was just 1.05 ± 0.21, a value statistically indistinguishable from 1 (Figures 3B, 4B and Tables S1, S2).

Additionally, to assess the reproducibility of the impact of the various components of hp3 in different mutant contexts, we calculated the fold effect on feedback regulation of deleting each of them by comparing multiple pairs of *rne-lacZ* reporters that differed by the presence or absence of a single hp3 component (Table S3). In each case, this was accomplished by dividing the fold effect (*ED*) in the presence of a given component by the corresponding fold effect in its absence. Gratifyingly, the impact of each component calculated in this manner proved to be context-independent and made clear their relative importance for feedback regulation: hp3D > hp3H ≈ hp3A ≈ hp3C > hp3B. Moreover, the product of multiplying their individual contributions (8.32) approximated their combined impact when hp3 was left entirely intact (7.22), suggesting a roughly additive effect of each component on feedback regulation.

In principle, hp3D might mediate feedback regulation by helping to recruit RNase E to the *rne* transcript. To test this possibility, we used an electrophoretic mobility shift assay to compare the ability of the catalytic domain of RNase E to bind hp3 in its entirety and a hp3 variant from which hp3D had been deleted (hp3ΔD). These experiments revealed that RNase E is able to bind hp3 with high affinity (Figure 5). Moreover, consistent with the regulatory importance of hp3D, the observed dissociation constant (*K*_d_) of intact hp3 was significantly lower than that of its mutant counterpart, a finding that suggests a possible role for RNase E binding in the mechanism by which hp3D contributes to the autoregulation of RNase E synthesis.

**Figure 5.**
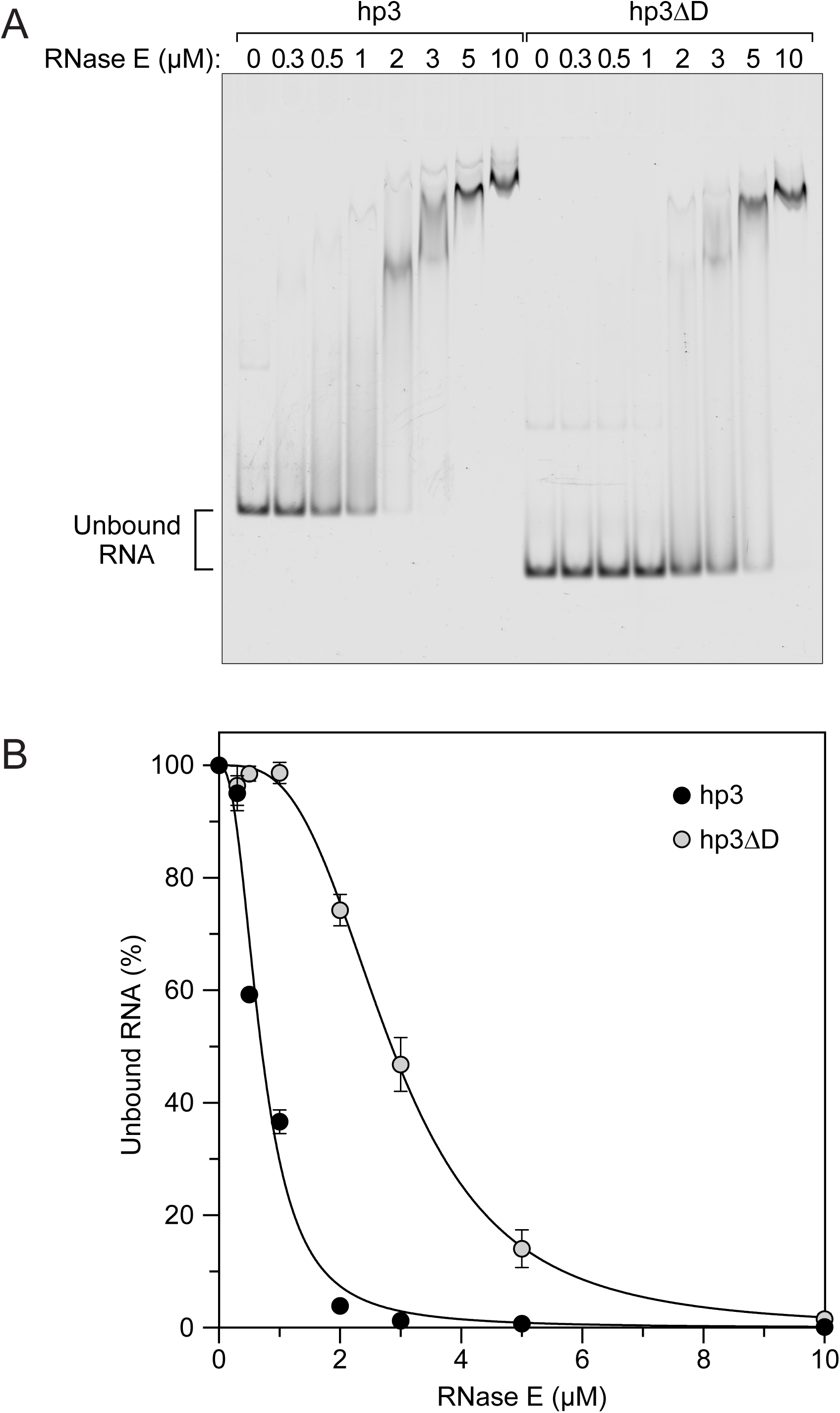
Effect of hp3D on the affinity of hp3 for the catalytic domain of RNase. **E.** (A) Electrophoretic mobility shift assays. Fluorescently labeled *rne* hp3 and hp3ΔD (Figure S2) were each combined with various concentrations of a catalytically inactive variant (D303A D346A) of the catalytic domain of RNase E (amino acids 1-529) and examined on a non-denaturing polyacrylamide gel for a change in electrophoretic mobility indicative of protein binding. (B) Graph. The percentage of unbound RNA was plotted as a function of RNase E concentration. Error bars (often not visible) each correspond to the standard deviation of multiple measurements. The observed dissociation constants were 0.70 ± 0.01 µM for hp3 and 2.85 ± 0.01 µM for hp3ΔD, a four-fold difference. These apparent *K*_d_ values undoubtedly were elevated by the inclusion of excess tRNA in the assay mixtures to suppress nonspecific binding.

## DISCUSSION

Previous research has shown that the ability of RNase E to autoregulate its synthesis in *E. coli* is controlled in cis by the *rne* 5′ UTR and that the upper portion of the second stem-loop in this 5′ UTR (hp2), comprising an apical loop and internal loop separated by two base pairs, makes an important contribution to RNase E autoregulation (16,17). We have now determined that another base-paired structure in the *rne* 5′ UTR, the branched stem-loop hp3, makes an even greater contribution to feedback regulation and that the impact of hp3 is mediated principally by a single one of its branches, hp3D, whose presence enables hp3 to bind the catalytic domain of RNase E with high affinity. The regulatory effects of hp2 and hp3 and of the various components of hp3 appear to be roughly additive.

The overall secondary structure of the *rne* 5′ UTR is well conserved in Enterobacterales despite considerable species-to-species sequence variation (Figure 1). Together with the translation initiation region, the components of this 5′ UTR with the greatest sequence conservation are the two now shown to control feedback regulation: hp3D and the top of hp2 (Figure S1), a finding consistent with their biological importance. In keeping with this observation, the hp3 component whose sequence and secondary structure are most variable, hp3B, has no detectable regulatory effect.

In view of previous *in vitro* evidence that RNase E binds to the top of hp2 (18) and our new evidence that it also binds to hp3 with a dissociation constant that is hp3D-dependent, we propose that these two 5′ UTR components recruit RNase E to *rne* mRNA by increasing the affinity of this endonuclease for the *rne* transcript. The additive regulatory effects of hp2 and hp3 suggest that they may bind independently to RNase E by interacting with distinct sites on the surface of this large, homotetrameric enzyme. A recruitment mechanism of this kind could enable autoregulation by sensitizing *rne* mRNA to the cellular concentration of its protein product.

This hypothesis is consonant with the critical importance of the internal loops of hp3D for feedback regulation, as RNase E has a strong affinity for single-stranded regions of RNA, sometimes in conjunction with a nearby base-paired structure (6,8,19,10,20,21). Moreover, the observation that RNase E appears to bind unpaired regions of RNA with little sequence specificity (2,3,21) is consistent with the substantial sequence variation observed for those two internal loops, which in Enterobacterales are less well conserved at the sequence level than both the base-paired regions of hp3D surrounding them and the top of hp2 (Figure S1). These observations suggest that the size, shape, and/or spatial positioning of the internal loops and adjacent base-paired regions of hp3D may be more important than the sequence of those internal loops for the regulatory function of hp3D. If so, the role of hp3C, the large internal loop of hp3A, and the four-way junction (hp3H), which each augment the impact of hp3D, may in part be to facilitate the interaction of RNase E with hp3D by influencing the location or orientation of that stem-loop and not simply to interact directly with RNase E themselves.

## MATERIALS AND METHODS

### Strains and plasmids

The isogenic *E. coli* strains CH1827 (MC1061, *rne*^+^ *zce-726::*Tn*10*) and CH1828 (MC1061, *rne-1 zce-726::*Tn*10*) are derivatives of MC1061 (F^-^*araD39* Δ(*ara-leu*)7697 Δ*lacX74 galE15 galK16 hsdR2 rpsL* (Str^R^)) (22). CH1828 carries a chromosomal *rne* missense mutation (*rne-1*, also known as *ams-1*) that reduces cellular RNase E activity at 37°C and eliminates it at 42°C {Jain, 1995 #227}. Plasmid pRNE101 is a pACYC177 derivative bearing the wild-type *E. coli rne* gene; the presence of this multicopy plasmid in CH1827 causes RNase E to be overproduced at ∼2.8 times its normal cellular concentration (15).

Plasmid pEZ1200 is a derivative of plasmid pEZ1000 (16) that contains an *rne-lacZ* reporter and confers ampicillin resistance. It has four modifications that distinguish it from its parent: (i) The segment of the *rne-lacZ* reporter corresponding to the *rne* coding region was reduced to just the first 29 codons, which were followed by a *Hin*dIII restriction site at the boundary between the *rne* and *lacZ* coding regions fused in-frame. (ii) An unnatural base pair inserted into pEZ1000 to create a BsrGI site in the *rne* 5′ UTR was removed to restore the wild-type 5′ UTR sequence. (iii) The *Hin*dIII and *Sal*I restriction sites 68-89 bp downstream of the 3′ end of the *lacZ* coding region were eliminated by removing 19 base pairs there (GCTTATCGATACCGTCGAC). (iv) Silent point mutations were incorporated into the first and second glycine codons of the *lacZ* coding region (both changed from GGG to GGA). Derivatives of pEZ1200 lacking specific elements of the *rne* 5′ UTR were constructed by overlap-extension PCR and restriction fragment substitution and verified by DNA sequencing. All *rne* 5′ UTR mutants lacking components of *rne* hp3 were examined in derivatives of plasmid pEZ1202, itself a pEZ1200 derivative encoding an *rne-lacZ* reporter in which hp3 remained intact but hp2 had been truncated. Plasmid pEZ1203 is a derivative of pEZ1200 encoding an *rne-lacZ* reporter in which hp2 remained intact but hp3 had been severely truncated so that only a small, unbranched stem-loop remained in its place. Plasmid pEZ1223 is a derivative of pEZ1200 encoding an *rne-lacZ* reporter in which both hp2 and hp3 had been truncated.

### β-galactosidase assays

Cultures of *E. coli* containing a derivative of plasmid pEZ1200 and either a high level of RNase E activity (CH1827 + pRNE101) or a low level of RNase E activity (CH1828) were grown overnight at 37°C or 30°C, respectively. The overnight cultures were diluted 1:1000 in triplicate into fresh LB medium (3 ml each) and grown with agitation at 37°C to log phase (OD_650_ = 0.2-0.8). Chloramphenicol was then added to each culture at a final concentration of 100µg/ml to stop further protein synthesis, followed by glycerol at a final concentration of 20% (v/v) to allow the cultures to be stored overnight at –80°C.

On a following day, the cells were thawed on ice, and β-galactosidase was assayed by colorimetrically monitoring *o*-nitrophenyl-β-D-galactopyranoside (ONPG) hydrolysis, as previously described (23). The OD_650_ values used to calculate β-galactosidase activity were those measured the previous day before adding glycerol and freezing the cells. Repression ratios were determined from nine pairwise comparisons by dividing the β-galactosidase activity in triplicate cultures containing low RNase E activity (CH1828) by the β-galactosidase activity in triplicate cultures containing high RNase E activity (CH1827 + pRNE101). The mean repression ratio (*R*) was then calculated from the resulting set of nine ratios, and the standard deviation of the ratios was calculated as the error. Repression ratios measured by this two-day procedure were nearly identical to those measured by the usual one-day procedure.

### Calculation of the fold effect of *rne* 5′ UTR mutations

To enable data from experiments performed on different days to be accurately compared, repression ratios were measured for the progenitor pEZ1202 reporter and the pEZ1223 reporter each time that a group of hp3 mutants was tested. The fold effect (*ED*) of each deletion within hp3 was calculated by dividing the repression ratio measured on the same day for the progenitor *rne-lacZ* reporter (encoded by pEZ1202) in which hp3 remained intact by the repression ratio of an otherwise identical reporter lacking one or more hp3 components of interest, after first subtracting 1 from each repression ratio.

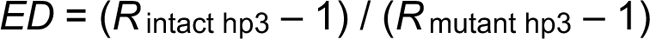

Conversely, to quantify the sufficiency of hp3 and its components for feedback regulation (*ER*), the repression ratio of intact hp3 or a mutant thereof that retained one or more hp3 components of interest and no others was divided by the repression ratio of an otherwise identical variant from which hp3 had been deleted altogether (Δhp3, encoded by pEZ1223), after first subtracting 1 from each repression ratio.

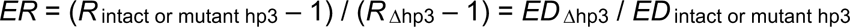

### Electrophoretic mobility shift assays

A catalytically inactive variant (D303A D346A) of the amino-terminal domain of *E. coli* RNase E (amino acids 1-529) bearing a carboxy-terminal hexahistidine tag was produced from plasmid pNRNE7529-D303A D346A in *E. coli* strain BL21(DE3) *rne*-131 Δ*rna* and purified by affinity chromatography on TALON beads (Takara Bio), as previously described (21). RNA ligands were synthesized for 8 hr at 37°C in a 100-μl reaction mixture containing 40 mM Tris-HCl (pH 7.9), 6 mM MgCl^2^, 10 mM NaCl, 2 mM spermidine, 10 mM dithiothreitol, 0.3 units/μl rRNasin (Promega), 1 mM ATP, 1 mM GTP, 1 mM CTP, 1 mM UTP, 10-50 ng/μl PCR-amplified DNA template (Table S4), and 5 units/μl T7 RNA polymerase (New England Biolabs). The resulting transcripts were treated with Turbo DNase (2 units/μl; Invitrogen) for 1 hr at 37°C in the buffer supplied by the manufacturer and then purified by gel electrophoresis and elution of the band of interest. The 5′-terminal segment of each RNA transcript was annealed to Cy5-Oligo20 (Cy5-GCGCTGCCGGGTCGCGTGGC) at a 2:1 molar ratio (0.6 μM RNA + 0.3 µM oligonucleotide) in 100 μl of 25 mM Bis-tris propane (pH 8.0), 100 mM NaCl by heating to 70°C for 10 min and then 50°C for 60 min, followed by slow cooling to <30°C. The resulting RNA-DNA duplexes were purified by nondenaturing polyacrylamide gel electrophoresis and elution.

RNA binding by RNase E was examined by electrophoretic mobility shift. Various concentrations (0-10 μM) of catalytically inactive RNase E (amino acids 1-529) were incubated for 30 min with a Cy5-labeled RNA-DNA duplex (∼27 nM) in a buffer containing 20 mM Tris chloride (pH 7.5), 50 mM potassium chloride, 5 mM magnesium chloride, 100 µg/ml *E. coli* tRNA, 1 mM dithiothreitol, 0.3 units/µl rRNasin, and 10% glycerol (v/v). Each mixture was then subjected to electrophoresis on a nondenaturing 4% polyacrylamide gel (29:1 acrylamide/bisacrylamide) in 1x TBE buffer. Changes in the fluorescence intensity of the band corresponding to unbound RNA were quantified by using a Typhoon RGB imager (Cytiva) and ImageQuant TL software (Cytiva) and used to calculate dissociation constants (*K*_d_) by fitting the data to a sigmoidal curve (Origin software) and determining the protein concentration at which half of the RNA remained unbound.

## SUPPLEMENTAL MATERIAL

Supplemental material (Figures S1 to S2 and Tables S1 to S4) is available for this article.

## ACKNOWLEDGMENTS

We thank Jamie Richards for his guidance and advice. This research was supported by grants to J.G.B. (R35GM145359 and R01GM035769) from the National Institutes of Health.

## AUTHOR CONTRIBUTIONS

J.G.B. conceived the project, S.E.E., T.B., and J.G.B. designed the experiments, S.E.E. and T.B. conducted the experiments, S.E.E., T.B., and J.G.B. interpreted the data, and J.G.B. wrote the paper.

