## Supplementary material for "Autoregulation of RNase E synthesis: Predominant impact of a branched RNA stem-loop to which the nuclease binds": Figures S1 to S2 and Tables S1 to S4

bioRχiv

This PDF file includes:

Figures S1 to S2
Tables S1 to S4

```
Escherichia       ---GUUUCCGUGUCCAUCCUUGUUAAAACAAGAAAUUUUACGGAAUAACCCAUU-UUGCC
Pantoea           --GUUUCUUACCUGAA--CCAUUUCAGGGCAGAAAAAGAGCGAAAUUGUCA-UU-UUGCC
Pectobacterium    GCGUUUCCCACGCGGAUUUCCGAUAAACAAGGGGAAAUUGCGAAGUUACUAAAU-UUGCC
Serratia          --ACUUUCCGCGUUGAUUCUCGUUAAA-CAGGGAAAACGGUGGAAUUAUAAAAU-UUGUC
Yersinia          ---AAUUCUACGUUGAUUCUCGUUAGACCAAGAGA-CUUGUGGAAUAAUAAACU-UUGCC
Hafnia            --GAUUUCCGCGUUAAUUCUCAUUAAGACAGGGAAAACGACGGGAUU-UGACGU-UUACU
Providencia       -------CUGCCAUAAUGUUUGAUAAAACGAUAAUUGGCAUGAAAUACAGAUUUGCUAUU
Budvicia          -GAUUUUCUGCCUUGAACCCGCCUAAUGCGUGGUAACAGGCAAAA-AAUAAAAU-UUG-C
                                 *       *                             *  *

Escherichia       CGACCGAUCAUCCA-CGCAGCAAUGGCGUAAGACGUAUUGAUCUUUCAGGCAGUUAGCGG
Pantoea           GGAUUGAGCAGACA-CGCAGCAAUGGCGUAAGACGUAAUGCGAAAUCAGGCACUUAGCGG
Pectobacterium    UGAUAACGCACACA-CGCAGCAAUGGCGUAAGACGUAAUGCGAAAUCAAGCAGUUAGCGG
Serratia          UGAUCGCGCAAAAA-CGCAGCAAUGGCGUAAGACGUAAUGCGAAAUCAAACAAUUAGCGG
Yersinia          UGAUCACACAAAAA-CGCGGCAAUGGCGUAAGACGUAAUGUGAAAUCAAGCAAUUAGCGG
Hafnia            UGAUGCCACAGAAA-UGCAGCAAUGGCGUAAGACGUAGUGUGAAAUCAAGUAAUUAGCGG
Providencia       UGAUUGAGUAAAAG-CGCGGCAAUGGCGUAAGACGCUUCGCAAAAUCAAAUAAUUAGCGA
Budvicia          AGAUCGCUCAGUCAGUGCAGCAAUGGCGUAAGACACCAUGAGAUAUCAAGCAAUUAGCGA
                   **      *      ** ***************     *     ***   * ******

Escherichia       GCUGCGGGUUGCAGUCCUUACCGGUAGAUGGAAAUA---------------UUUCUGGAG
Pantoea           GCUGCGGGUUGCGGCCUGGACAACAGUAUGGGAUGCAC-----------GGGUUUUCUUU
Pectobacterium    GCUGUGGAUUGCAGCUUGGCCGGCAA-AUGGAAUCAGA-------------UCUGGCGAC
Serratia          GCUGCGGGUUGCAGCUUGGCCGGCAA-AUGGAAUCAGA-------------UCUGUCGAC
Yersinia          GCUGCGGGUUGCAGCUUGGCCGGCAA-GUGGGUAAAGC-------------UCUGUUGUC
Hafnia            GCUGUGGGUAGCAGUCUGACCUAAAACAUGGAAUCAGCAAUGCUGCAUAUUUUUGCAAUU
Providencia       GCAACGAAUAUUAGCUUUUUGGGUUA---GGAACUUCC-------------GUUAACAAA
Budvicia          GCUACGGGUAGUGGCUUGGAUGGUAA-AAGGAACCCGA-----------UCUCUAUCUAA
                  **   *  *    *               **                      *

Escherichia       AGUAAUACCCAGUCUGUUUCUUUGAUAAUUGCGCUGU-UUUUCCGCAUGAAAAACGGGC-
Pantoea           AUUCUCUCAGGCAUU-UCCACUAUCU---GGUGCUGU-UUUCCCUUAUGAAAAACAGGC-
Pectobacterium    AUUAU-UCAGAAGCUGUUCCCUCAGUAAAUGCGCUGU-UUUCCAUCAGGAAAUACAGGC-
Serratia          GUAAA-UCAGAGGCUAUUCCCCUAGUAAAAGUGCUGU-CUUUCACAAAGAAAAGCAGGCU
Yersinia          UUAAAUUCAGAAGCUACCCUUUAUAUAAAAGCGCUGU-UUUUCAGAACGAAACACAGGCU
Hafnia            AUUGACUGAUAAACAGUGCUGUGUUUCACAAAGCUGUAUUUUCGCUAAGAAACAUAGGCU
Providencia       AUUAU------GACGGCUUCCUGAUUAAGAACGCUGAAUUUUUAAUCUAAAAAUCAGGUU
Budvicia          CUUUAGCGACGGACU-UUCCCUGAUAAAAAGUGCUG--UUUUCACAAAGAAAUACAGACU
                                                  ****   **        ***    *

E_coli            AACCGACACU--CUGCGCCUCUUUGAGCUGACGAUAACCGUGAGGUUGGCGAC---GCGA
Pantoea           CACCGACGUUAUUUGCGCCCCUCGCG-CGGUCGACAGCCGUGAGGUUGACGACAUUGCAC
Pectobacterium    UACCGAAAC---AUGCGUCUCUAU--GCAGGCGACAACCGGGAGGUUGACGUCCCUGCGA
Serratia          UACCGAGAUA--AUGCGCCCCUAU--GCAGGCGACAACCGUGAGGUUGACGACUUUGCGA
Yersinia          UACCGAAAUA--AUGCGCCCCUAU--GCAGGCGACAACCGUGAGGUUGACGGCUUAGCUA
Hafnia            UACCGAAAU---AUGCGCCCCUAU--GCAGGCGACAACCGUGAGGUUGACGACUUUGCGA
Providencia       CACCGA-AUA--CUGCGCGUCUCUAUACGGACGACUGCCGGGAGGCAGGCGGCUUUGUAA
Budvicia          UACCGAGAUA--AUGCGUCCCUGU--GCAAACGACAACCGUGAGGUUGACGAUUUUGCGA
                   *****       ****   **     *   ***   ***  ****  *  **     *

E_coli            CUAGACACGAGGCCAUCGGUUCA-CACCCGGAAAGGCG--UUACUUUGCCCGCAG-CUUA
Pantoea           UAAGUCACGGGGCCAUCGGUUGA-UACUCGUCCAGGG---UAACUAUGCCCGCAGCUGUA
Pectobacterium    UAAGCCACGAGGCCAUCGGUUCA-CUC-CGGUCAUGCGGUUCUUUUUGUCCGCAG-CUCU
Serratia          GAAGACACGGGGCUUUCGGUUUA-ACUCCGGCCAUGAGG-UUAUUUUGCCCGCAG-CUUU
Yersinia          GAAGACUCGGGGCCAUCGGUUUA---UCCGGUCAUGAGG-UUAUUUUGCCCGCAG-UUCU
Hafnia            UAAGACUCGGGGUCAUCGGUCUUCCUCUCGGUCAGGACG-UUAUUUUGCCCGCAG-CUUA
Providencia       -AAAUCACGAGGCCAUCGGUUUA-CAAUAGCUC-------CCUUUUUCGCUGCU--CUUU
Budvicia          UAAGACUCGGGGCCAUCGGUCAA-CUC-CGUCCAUGAG--UAUUCUUAUACGUAGACUCA
                    *  * ** **   *****         *                *    *
```

```
E_coli              GUCGUCAAU----GUAAGAAUAAUGAGUAAGUU--ACGAUG
Pantoea             AUCACUCAU----GAAAGAAUAACGAGUAAGUC--ACGAUG
Pectobacterium      AUCAAUAAUGCAAGUAAAAAUAACGAGUAAGUUG-AAGAUG
Serratia            GUCGAUAAU----GUAAAAAUAACGAGUAAGUU--AAGAUG
Yersinia            AACGCCAAU----GUAAAAAUAACGAGUAAGUUA-AAGAUG
Hafnia              GUUGCCAAU----GCAAAAAAAAUGAGUAAGUUG-AAGAUG
Providencia         AUUGAAAGU-----AAAAAAUAAUGAGUAAGUAACAAUAUG
Budvicia            AUUGACAAU----GUAAAAA-AAUGAGUAAAGUU-UAAAUG
                            *      ** ** ** ******       ***
```

**Figure S1. Sequence alignment of the *rne* 5′ untranslated region in Enterobacterales.** The aligned 5′ UTR sequences of *rne* transcripts representing the Enterobacterales families Enterobacteriaceae (*Escherichia coli*), Erwiniaceae (*Pantoea agglomerans*), Pectobacteriaceae (*Pectobacterium atrosepticum*), Yersiniaceae (*Serratia marcescens* and *Yersinia pestis*), Hafniaceae (*Hafnia alvei*), Morganellaceae (*Providencia stuartii*), and Budviciaceae (*Budvicia aquatica*) are shown. The two 5′ UTR components with the greatest impact on feedback regulation are colored green (hp2) or red (hp3D), with a blue background highlighting the nucleotides that correspond to the internal loops hp3Da and hp3Db. Strictly conserved nucleotides are identified with an asterisk (*), and the Shine-Dalgarno element and initiation codon of each transcript are underlined.

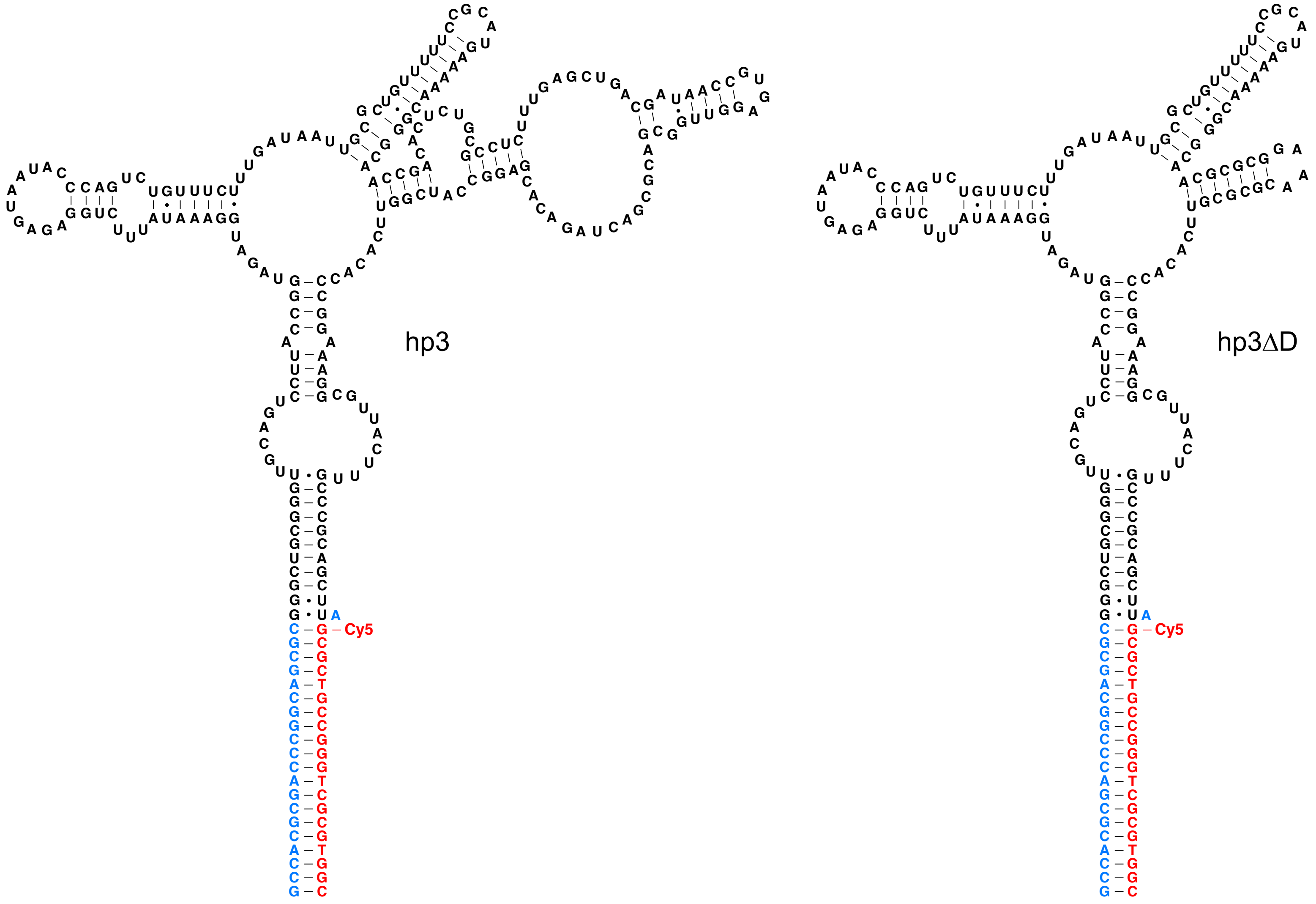

**Figure S2. Fluorescently labeled RNA ligands used in RNase E binding assays.** Ribonucleotides corresponding to hp3 and hp3ΔD are colored black. Additional ribonucleotides in each RNA ligand are colored blue. A complementary deoxyoligonucleotide 5′-terminally labeled with Cy5 is colored red.

**Table S1. Effect of deleting *rne* hp3 components**

| Deleted hp3 component [a] | Fold effect (*ED*) vs pEZ1202 [b] |
|---|---|
| Δhp3A | 1.53 ± 0.39 |
| Δhp3B | 1.01 ± 0.25 |
| Δhp3C | 1.30 ± 0.28 |
| Δhp3D | 3.12 ± 0.71 |
| Δhp3Dab | 3.79 ± 0.76 |
| Δhp3Da | 2.04 ± 0.49 |
| Δhp3Db | 2.63 ± 0.53 |
| Δhp3ABC | 1.75 ± 0.34 |
| Δhp3ABCH | 2.42 ± 0.47 |
| Δhp3BC | 1.36 ± 0.25 |
| Δhp3BCH | 2.02 ± 0.43 |
| Δhp3BCD | 3.97 ± 0.69 |
| Δhp3BCDH | 6.85 ± 1.22 |

[a] The effect of deleting hp3 components was determined in *rne-lacZ* reporters lacking hp2.

[b] Plasmid pEZ1202 encodes an *rne-lacZ* reporter in which hp3 remains intact but hp2 has been deleted.

**Table S2. Effect of retaining *rne* 5′ UTR components**

| Retained 5′ UTR component [a] | Fold effect (*ER*) vs pEZ1223 [b] |
|---|---|
| hp2 | 3.78 ± 0.41 |
| hp3 | 7.22 ± 0.67 |
| hp3ADH | 5.30 ± 1.08 |
| hp3AD | 3.58 ± 0.83 |
| hp3D | 2.98 ± 0.64 |
| hp3ABCH | 2.32 ± 0.57 |
| hp3AH | 1.82 ± 0.36 |
| hp3A | 1.05 ± 0.21 |

[a] The effect of retaining hp2 was determined by analyzing a pEZ1203-encoded *rne*-*lacZ* reporter in which hp2 remained intact but hp3 had been deleted. The effect of retaining hp3 or only some of its components was determined in *rne-lacZ* reporters lacking hp2. By definition, hp3 = hp3ABCDH.

[b] Plasmid pEZ1223 encodes an *rne-lacZ* reporter lacking both hp2 and hp3.

**Table S3. Effect of deleting hp3 components in disparate structural contexts**

| hp3 component [a] | Mutant comparison 1 | Fold effect (*ED*) 1 | Mutant comparison 2 | Fold effect (*ED*) 2 | Mean |
|---|---|---|---|---|---|
| hp3A | Δhp3A / Intact hp3 | 1.53 ± 0.39 | Δhp3ABC / Δhp3BC | 1.28 ± 0.34 | 1.41 ± 0.21 |
| hp3B | Δhp3B / Intact hp3 | 1.01 ± 0.25 | Δhp3BC / Δhp3C | 1.05 ± 0.30 | 1.03 ± 0.15 |
| hp3C | Δhp3C / Intact hp3 | 1.30 ± 0.28 | Δhp3BC / Δhp3B | 1.35 ± 0.41 | 1.33 ± 0.19 |
| hp3D | Δhp3D / Intact hp3 | 3.12 ± 0.71 | Δhp3BCD / Δhp3BC | 2.91 ± 0.74 | 3.02 ± 0.39 |
| hp3H | Δhp3BCH / Δhp3BC | 1.48 ± 0.42 | Δhp3ABCH / Δhp3ABC | 1.39 ± 0.38 | 1.44 ± 0.57 |

[a] The effect of deleting each hp3 component was determined in *rne-lacZ* reporters lacking hp2.

**Table S4. Templates for *in vitro* transcription of RNase E ligands**

| RNA | DNA template [a] |
|---|---|
| hp3 | AAATT**TAATACGACTCACTATA**GCCACGCGACCCGGCAGCGCGGGCTGCGGGTTGCAGTCCTTACCGGTAGATGGAAAT<br>ATTTCTGGAGAGTAATACCCAGTCTGTTTCTTTGATAATTGCGCTGTTTTTCCGCATGAAAAACGGGCAACCGACACTCTG<br>CGCCTCTTTGAGCTGACGATAACCGTGAGGTTGGCGACGCGACTAGACACGAGGCCATCGGTTCACACCCGGAAAGGC<br>GTTACTTTGCCCGCAGCTTA |
| hp3ΔD | AAATT**TAATACGACTCACTATA**GCCACGCGACCCGGCAGCGCGGGCTGCGGGTTGCAGTCCTTACCGGTAGATGGAAAT<br>ATTTCTGGAGAGTAATACCCAGTCTGTTTCTTTGATAATTGCGCTGTTTTTCCGCATGAAAAACGGGCAACGCGCGGAAAC<br>GCGCGTTCACACCCGGAAAGGCGTTACTTTGCCCGCAGCTTA |

[a] The T7 promoter is shown in boldface. Additional transcribed nucleotides that were added to hp3 to create a binding site for a Cy5-labeled DNA oligonucleotide (Figure S2) are shown in blue.
